# p53 induces ARTS to promote mitochondrial apoptosis

**DOI:** 10.1101/2020.05.14.096982

**Authors:** Qian Hao, Jiaxiang Chen, Junming Liao, Yingdan Huang, Sarit Larisch, Shelya X Zeng, Hua Lu, Xiang Zhou

## Abstract

<u>A</u>poptosis <u>R</u>elated protein in <u>T</u>GF-β <u>S</u>ignaling pathway (ARTS) was originally discovered in cells undergoing apoptosis in response to TGF-β, but ARTS also acts downstream of many other apoptotic stimuli. ARTS induces apoptosis by antagonizing the anti-apoptotic proteins XIAP and Bcl-2. Here, we identified the pro-apoptotic *Sept4/ARTS* gene as a p53-responsive target gene. Ectopic p53 and a variety of p53-inducing agents increased both mRNA and protein levels of ARTS, whereas ablation of p53 reduced ARTS expression in response to multiple stress conditions. Also, γ-irradiation induced p53-dependent ARTS expression in mice. Consistently, p53 binds to the responsive DNA element on the ARTS promoter and transcriptionally activated the promoter-driven expression of a luciferase reporter gene. Interestingly, ARTS binds to and sequesters p53 at mitochondria, enhancing the interaction of the latter with Bcl-XL. Ectopic ARTS markedly augments DNA damage stress- or Nutlin-3-triggered apoptosis, while ablation of ARTS preferentially impairs p53-induced apoptosis. Altogether, these findings demonstrate that ARTS collaborates with p53 in mitochondria-engaged apoptosis.

## Introduction

The tumor suppressor p53 prevents genomic instability and tumorigenesis through multiple mechanisms. As a transcription factor, it arrests cell cycle progression and facilitates DNA repair by inducing the RNA expression of a number of cell cycle- and DNA repair-associated genes when pre-cancerous cells undergo mild insult of replication stress. In addition, p53 can elicit cell death through upregulation of the pro-apoptotic genes, such as BAX, NOXA and PUMA, when cancer cells are subjected to severe DNA damage stress, such as therapeutic intervention (Riley et al., 2008, Levine, 2019). However, independently of its transcriptional activity, p53 has also been shown to promote apoptosis via mitochondria-involved mechanisms (Green and Kroemer, 2009). p53 was shown to bind to Bcl-XL via its DNA-binding domain (DBD) and de-represses the mitochondrial BAK, BAX and PUMA, consequently leading to mitochondrial outer membrane permeabilization (MOMP) and release of cytochrome c (Mihara et al., 2003, Moll et al., 2006, Leu et al., 2004). p53 is such detrimental to cancer cells that several auto-regulatory mechanisms have been evolved in cancer cells to control its activity (Zhou et al., 2017). The E3-ubiquitin ligase MDM2, encoded by a p53 target gene, is the master negative regulator that can inhibit p53 activity by directly concealing its transcriptional activation domain (TAD) and promoting its proteolytic degradation (Liu et al., 2019, Oliner et al., 1993, Haupt et al., 1997, Kubbutat et al., 1997). Other p53-inducible proteins, such as NGFR and PHLDB3 (Zhou et al., 2016, Chao et al., 2016), have been shown to either directly or collaborate with MDM2 to repress p53 as negative feedback regulators. This study as presented here identified ARTS (apoptosis-related protein in the TGF-β signaling pathway) as another p53 target that could play a role in regulation of p53’s apoptotic activity as well.

ARTS is a pro-apoptotic protein located at the outer membrane of the mitochondria (Larisch et al., 2000, Edison et al., 2012). ARTS protein is derived from alternative splicing of the *SEPT4* gene (Larisch et al., 2000, Mandel-Gutfreund et al., 2011). Although ARTS was originally discovered in cells induced for apoptosis by TGF-β, it was later found that ARTS acts downstream of basically all apoptosis stimuli tested, such as treatment with STS (staurosporine), etoposide, arabinoside (Ara-c), nocadosole, UV radiation, TNF-α, etc. (Larisch et al., 2000, Elhasid et al., 2004, Lotan et al., 2005). ARTS initiates caspase activation upstream of mitochondria by directly binding and degrading XIAP (X-linked inhibitor of apoptosis) via the ubiquitin proteasome system (UPS) (Gottfried et al., 2004, Garrison et al., 2011, Edison et al., 2012). Recently, ARTS was shown to induce ubiquitination and degradation of Bcl-2 by bridging the E3-ubiquitin ligase XIAP to Bcl-2 (Edison et al., 2017). Studies in human and mice have shown that ARTS functions as a tumor suppressor protein (Elhasid et al., 2004, Kissel et al., 2005, Garcia-Fernandez et al., 2010, Fuchs et al., 2013, Koren et al., 2018). Moreover, *Sept4/ARTS* deficient mice exhibit high levels of stem and progenitor cells which are resistant to apoptosis (Garcia-Fernandez et al., 2010, Fuchs et al., 2013, Koren et al., 2018).

When screening for novel p53 target genes by microarray analysis of Inauhzin-treated cancer cells (Liao et al., 2012) [of note, Inauhzin is a p53 activating small molecule identified by our lab (Zhang et al., 2012)], we identified ARTS as a potential p53 target gene. Our further study of this molecule not only confirmed that p53 transcriptionally induces ARTS expression in cancer cells and in mice, but also revealed that ARTS interplays with p53 in inhibition of Bcl-XL in the mitochondria, consequently augmenting p53-dependent apoptosis. As detailed below, our findings demonstrate that ARTS can enhance p53-mediated mitochondrial apoptosis in a positive feedback fashion.

## Materials and Methods

### Plasmids and antibodies

The Flag-tagged pcDNA-ARTS plasmid was generated by inserting the full-length ARTS cDNA amplified from the pcDNA3-Myc-ARTS plasmid as a gift from Dr. Sarit Larisch into the pcDNA3.0/Flag vector. The plasmid encoding Flag-Bcl-XL was purchased from OriGene Technologies (Rockville, MD, USA). The plasmids encoding p53, HA-MDM2, and His-Ub were described previously (Zhou et al., 2013). The pGL3-RE1, RE2, and RE3 plasmids were generated by inserting the genomic DNA covering p53 RE1, RE2 or RE3 into the pGL3-promoter vector using the following primers, 5’ -CGGGGTACCATTCAGCAGGTGCCAGGAA-3’ and 5’-CCGCTCGAGACGATACAGTCAGAGAGTCCTT-3’ for RE1; 5’ -CGGGGTACCGTATTAGACCCTGCCTCCATCA-3’ and 5’-CCGCTCGAGGAAGACTGACTTTGAGCCATCC-3’ for RE2; 5’ -CGGGGTACCTGCCTCGGACTCCTGAGTA-3’ and 5’-CCGCTCGAGGGGACAGACAAGCAGAGAAAC-3’ for RE3. The lentivirus-based ARTS-overexpressing or shRNA plasmid was constructed using the vectors pLenti-EF1a-EGFP-P2A-Puro-CMV-3Flag and pLKD-CMV-G…PR-U6, respectively (OBio Technology, Shanghai, China). The shRNA sequence targeting ARTS was previously described (Edison et al., 2012). The anti-ARTS (Sigma-Aldrich, St louis, MO, USA, monoclonal mouse cat # A4471), anti-p53 (Santa Cruz Biotechnology, Santa Cruz, CA, USA), anti-Flag (Sigma-Aldrich), anti-HA (Cell Signaling Technology, Danvers, MA, USA), anti-p21 (Cell Signaling Technology), anti-PUMA (Cell Signaling Technology), anti-COX IV (Cell Signaling Technology), anti-GAPDH (Proteintech, Wuhan, Hubei, China), and anti-β-actin (Proteintech) were commercially purchased.

### Cell culture and transient transfection

Human cancer cell lines HCT116^p53+/+^, HCT116^p53-/-^, H460, H1299, SK-MEL-103, SK-MEL-147 were cultured in Dulbecco’s modified Eagle’s medium (DMEM) supplemented with 10% fetal bovine serum, 50 U/ml penicillin and 0.1 mg/ml streptomycin. All cells were maintained at 37°C in a 5% CO^2^ humidified atmosphere. Cells seeded on the plate overnight were transfected with plasmids or siRNA as indicated in the figure legends using Hieff TransTM Liposomal transfection reagent following the manufacturer’s protocol (Yeasen, Shanghai, China). Cells were harvested at 30-72 hrs post-transfection for designed experiments. The proteasome inhibitor MG132 was purchased from Sigma-Aldrich.

### RNA interference

siRNA against p53 was commercially purchased (GenePharma, Shanghai, China). The amount of 40 to 100 nM of siRNA was introduced into cells using Hieff TransTM Liposomal transfection reagent following the manufacturer’s protocol. Cells were harvested 48 to 72 hrs after transfection for IB or RT-qPCR. The sequence of the siRNA used here was GUAAUCUACUGGGACGGAA and as previously described (Zhou et al., 2016).

### Reverse transcription and quantitative real-time PCR

RNA was isolated from cells using RNAiso Plus following the manufacturer’s protocol (Takara, Dalian, Liaoning, China). Total RNAs of 0.5 to 1 μg were used as templates for the reverse transcription using PrimeScriptTM RT Reagent Kit with gDNA Eraser (Takara). Quantitative PCR (qPCR) was conducted using TB GreenTM Premix Ex TaqTM (Tli RNaseH Plus) according to the manufacturer’s protocol (Takara). The primers used for qPCR were 5’-ACCATTGTGGACACACCAGG-3’ and 5’-GAACCTGTGACCACCTGCTA-3’ for human ARTS, 5’-CAGGGCAGGGCTACCACTAG-3’ and 5’-TGATGCAGGGCCTTCATGA-3’ for mouse ARTS. The primers for human and mouse p21 were previously described (Zhou et al., 2015, Zhang et al., 2017).

### Immunoblotting

Cells were harvested and lysed in lysis buffer consisting of 50 mM Tris/HCl (pH7.5), 0.5% Nonidet P-40 (NP-40), 1 mM EDTA, 150 mM NaCl, 1 mM dithiothreitol (DTT), 0.2 mM phenylmethylsulfonyl fluoride (PMSF), 10 μM pepstatin A and 1 μg/ml leupeptin. Equal amounts of clear cell lysate (20-80 μg) were used for immunoblotting (IB) analysis as described previously (Zhou et al., 2013).

### Immunoprecipitation

Immunoprecipitation (IP) was conducted using antibodies as indicated in figure legends. Briefly, 500 to 1000 μg of proteins were incubated with the indicated antibody at 4°C for 4 hrs or overnight. Protein A or G beads (Santa Cruz Biotechnology) were then added, and the mixture was incubated at 4°C for additional 1 to 2 hr. Beads were washed at least three times with lysis buffer. Bound proteins were detected by IB with antibodies as indicated in figure legends.

### Luciferase reporter assay

Cells were transfected with pGL3-RE1, RE2 or RE3 plasmid together with pGMLR-TK, and the p53-encoding plasmid or the pcDNA vector as indicated in the figure. Dual-Luciferase Reporter Assay System was used to determine luciferase activity according to the manufacturer’s instruction (Promega, Madison, WI, USA).

### Chromatin immunoprecipitation

Chromatin immunoprecipitation (ChIP) assay was performed using antibodies as indicated in figure legends and described previously (Liao and Lu, 2013). The reverse cross-linked immuoprecipitated DNA fragments were purified using GeneJET gel extraction kit (Thermo Scientific, Waltham, MA, USA) followed by PCR analysis for the p53-responsive DNA elements on the human ARTS promoter using the following primers, 5’-GTATTAGACCCTGCCTCCATCA-3’ and 5’-GAAGACTGACTTTGAGCCATCC-3’.

### Subcellular fractionation

Cells suspension in the fractionation buffer (20 mM HEPES (pH 7.4), 10 mM KCl, 2 mM MgCl_2_, 1 mM EDTA, 1 mM EGTA, 1 mM DTT and protease inhibitors) was incubated for 15 min on ice and went through a 27 gauge needle 10 times. After incubation on ice for another 20 min, the samples were centrifuged at 3,000 rpm for 5 min. The pellets contained nuclei. Supernatants were centrifuged at 8,000 rpm for 5 min. Pellets contained mitochondria and supernatants contained the cytoplasm and membrane fraction.

### γ-irradiation of mice

p53^+/+^ and p53^-/-^ mice of eight weeks of age were subjected to whole-body γ-irradiation (5 Gy) at a dose rate of 0.75 Gy/min. Mice were sacrificed and their spleens and thymuses were harvested 0 or 6 h post-irradiation (Bouvard et al., 2000). The tissues were analyzed by IB and RT-qPCR for p53, p21 and ARTS expression.

### Flow cytometry analysis

The PE Annexin V Apoptosis Detection Kit I (BD Biosciences, San Diego, CA, USA) was used for apoptosis analysis according to the manufacturer’s instruction. Briefly, cells were washed twice with cold PBS and then re-suspended in Annexin V Binding Buffer at a concentration of 1 x 10^6^ cells / ml. Cells were incubated with PE Annexin V and 7-AAD for 15 min at RT in the dark. Flow cytometry was performed using a FC500 MPL flow cytometer (Beckham coulter, Indianapolis, IN, USA) within 1 hr.

### Cell viability assay

The Cell Counting Kit-8 (CCK-8) (Dojindo Molecular Technologies, Japan) was used according to the manufacturer’s instructions. Cells of 2,000 to 5,000 were seeded per well in 96-well culture plates at 12 hr post-transfection. Cell viability was determined by adding WST-8 at a final concentration of 10% to each well, and the absorbance of the samples was measured at 450 nm using a Microplate Reader every 24 hr for 4-5 days.

### Statistics

The Student’s t-test or one way analysis of variance (ANOVA) was performed to evaluate the differences between two groups or more than two groups. p<0.05 was considered statistically significant. All the data are presented as mean ± SD.

## Results

### ARTS expression is induced by p53 in cancer cells

Through a primary screen for p53-responsive genes by the p53-inducing agent Inauhzin (INZ) (Zhang et al., 2012, Liao et al., 2012), we identified ARTS as a possible p53 target gene. To confirm this, we ectopically expressed p53 in H460 lung cancer cells, and indeed found that the expression of ARTS is elevated at both mRNA (Figure 1A) and protein levels (Figure 1B). Also, we showed that exogenous p53 induces ARTS mRNA (Figure 1C) and protein expression (Figure 1D) in HCT116^p53+/+^ colon cancer cells. To verify these observations, several p53-inducing agents were used to test if ARTS expression is responsive to p53 activation. As shown in Figures 1E and 1F, 5-fluorouracil (5-FU), Doxorubicin (DOX) or Inauhzin treatment dramatically stimulated both mRNA and protein expression of ARTS in H460 cells. Consistently, ARTS expression could be induced by 5-fluorouracil and Doxorubicin in two wild-type p53-harboring melanoma cell lines, SK-MEL-147 and SK-MEL-103 (Figures 1G and 1H). In addition, we found that ARTS expression can also be elevated upon oxidative stress triggered by H_2_O_2_ (Figure 1I) that was shown to activate p53 (Levine and Oren, 2009, Eriksson et al., 2019). The induction of ARTS expression observed could be specifically owing to p53 activation, given that the p53 target gene p21 was simultaneously induced under the same conditions (Figures 1A–1I) and that ectopic mutant p53-R175H exerted no effect on ARTS or p21 expression (Figure 1J). Then, we determined if endogenous p53 is required for ARTS induction upon different stress signals. ARTS mRNA levels were assessed in HCT116^p53+/+^ cells treated with Inauhzin, Cisplatin, 5-fluorouracil or Nutlin-3 following knockdown of p53. As expected, the expression of ARTS in response to these treatments was significantly reduced upon p53 depletion (Figure 1K), which is further validated by the protein expression of ARTS in H460 cells under both normal and DNA damage conditions (Figure 1L). Therefore, these results indicate that the *Sept4/ARTS* gene is a p53-inducibale gene in response to various stress signals in cancer cells.

**Figure 1.**
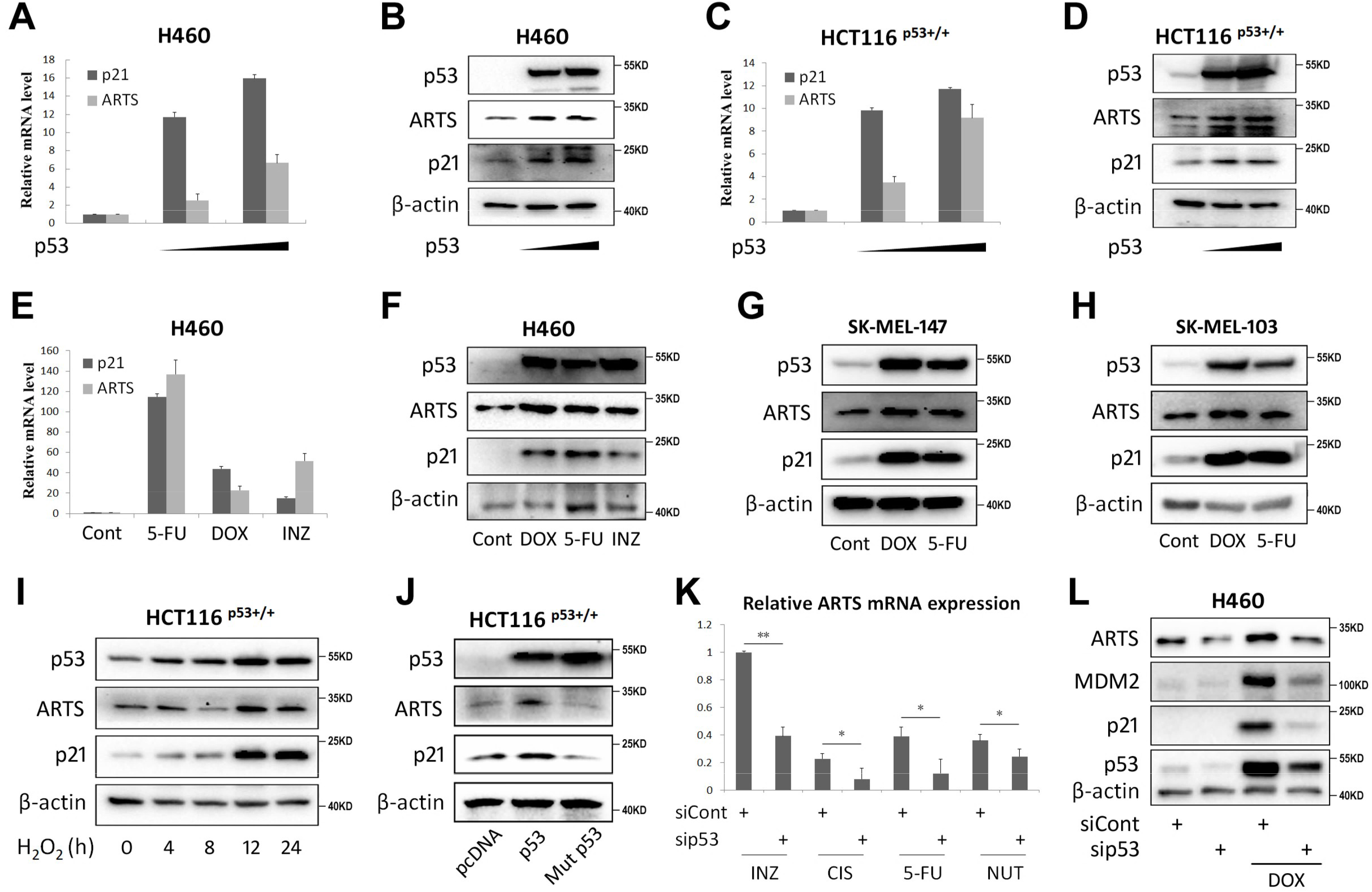
p53 induces ARTS expression in cancer cells. (A, B) Ectopic p53 induces ARTS mRNA (A) and protein (B) expression in H460 cells. Cells were transfected with the vector or increased doses of p53 plasmid followed by RT-qPCR or IB analysis. (C, D) Ectopic p53 induces ARTS mRNA (C) and protein (D) expression in HCT116^p53+/+^ cells. Cells were transfected with the vector or increased doses of p53 plasmid followed by RT-qPCR or IB analysis. (E, F) The p53-inducing agents elevate ARTS mRNA (E) and protein (F) levels in H460 cells. Cells were treated with 5-fluorouracil, Doxorubicin or Inauhzin for 18 h followed by RT-qPCR or IB analysis. (G, H) The p53-inducing agents elevate ARTS expression in melanoma cell lines. Cells were treated with 5-fluorouracil or Doxorubicin for 18 h followed by IB analysis. (I) H_2_O_2_-induced oxidative stress increases ARTS expression. Cells were treated with H_2_O_2_ for the indicated time followed by IB analysis. (J) Wild-type p53, but not mutant p53-R175H, induces ARTS expression. Cells were transfected with plasmids as indicated followed by IB analysis. (K) Ablation of p53 reduces ARTS mRNA expression upon multiple stress conditions. Cells were transfected with p53 siRNA and treated with agents as indicated 18 h before harvest for RT-qPCR analysis. (L) Ablation of p53 reduces ARTS protein expression upon DNA damage stress. Cells were transfected with p53 siRNA and treated with Doxorubicin 18 h before harvest for IB analysis.

### γ-irradiation induces ARTS expression dependent on p53 in mice

Since ARTS is required for tumor suppression *in vivo* (Garcia-Fernandez et al., 2010, Elhasid et al., 2004), we tested if this tumor suppressor can be activated through p53 in mice. The p53^+/+^ and p53^-/-^ mice were exposed to γ-irradiation, and the radiosensitive organs, thymuses and spleens (Bouvard et al., 2000), were harvested for analysis of the expression of murine p53, ARTS and p21. As shown in Figure 2A, γ-irradiation drastically boosted the protein levels of p53 and ARTS in the thymuses of the p53^+/+^ mice, but not in those of the p53^-/-^ mice. The induction of ARTS in response to γ-irradiation could be due to the increased transcriptional activity of p53, as evidenced by the upregulation of its mRNA level (Figure 2B). In line with these data, the irradiated spleens also displayed higher expression of murine ARTS in a p53-dependent fashion (Figures 2C and 2D). Together with the results in Figure 1, these findings demonstrate that p53 induces ARTS expression in response to various stressors not only in cancer cells but also in mice.

**Figure 2.**
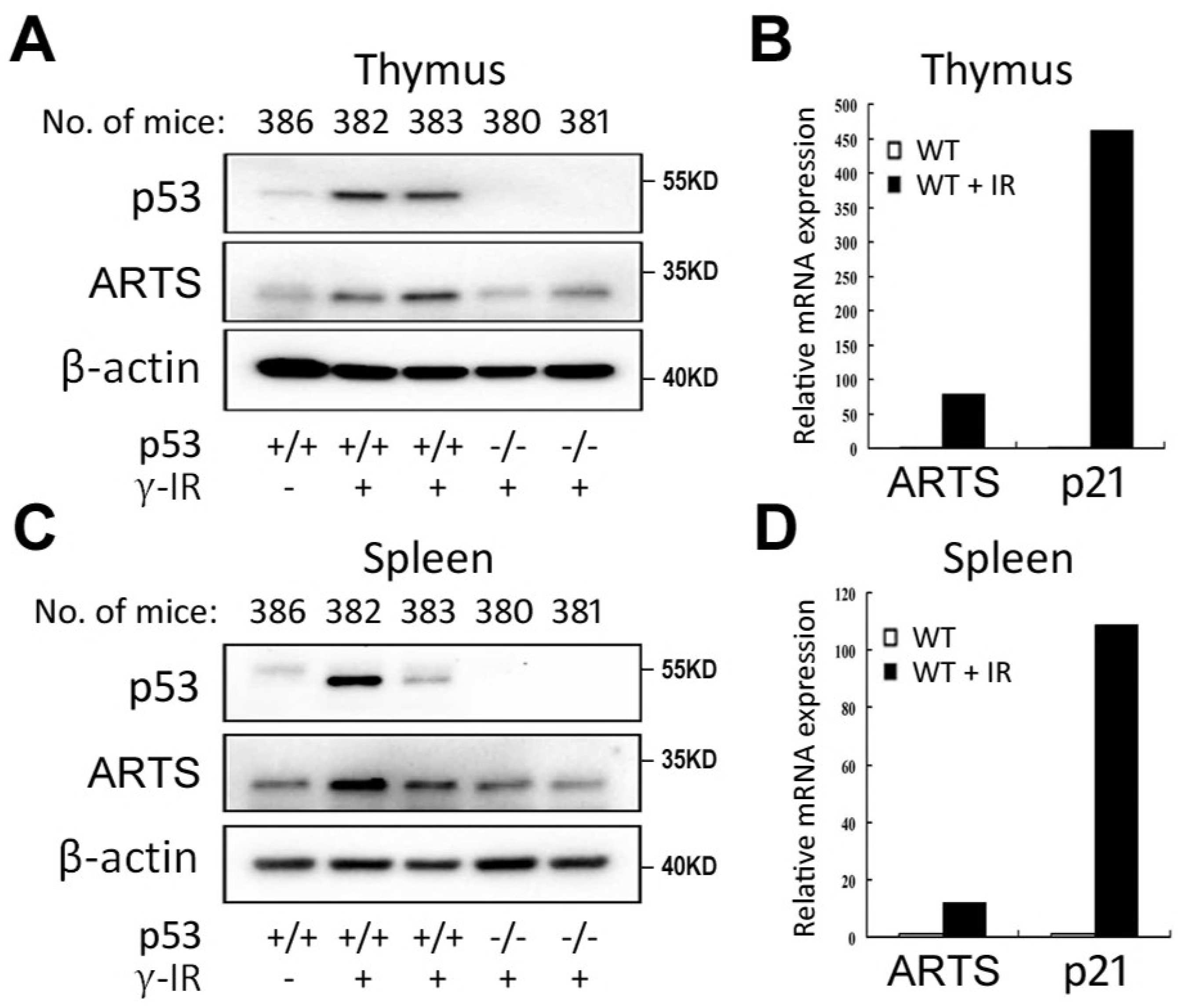
ARTS expression is induced by γ-irradiation through p53 in mice. (A, B) ARTS mRNA (A) and protein (B) expression is elevated in the irradiated murine thymuses. The irradiated mice were sacrificed, and the thymuses were freshly harvested and subjected to IB or RT-qPCR analysis. (C, D) ARTS mRNA (C) and protein (D) expression is elevated in the irradiated murine spleens. The irradiated mice were sacrificed, and the spleens were freshly harvested and subjected to IB or RT-qPCR analysis.

### p53 transcriptionally activates ARTS expression by associating with its promoter

Since p53 mainly functions as a transcription factor, we speculated that p53 may enhance *Sept4/ARTS* gene transcription by binding to its promoter. Indeed, by carefully analyzing the genomic sequence of the human *Sept4/ARTS* gene using p53MH algorism (Hoh et al., 2002), we found three potential p53-responsive elements (REs) located at −3087 and −2279 bp before the transcription-start site (TSS), and +9720 bp within the intron of the *Sept4/ARTS* gene (Figure 3A). To determine if p53 activates transcription of *ARTS* through any of these REs, we tested the luciferase reporter gene expression driven by each of the REs. Remarkably, p53 induced luciferase activity only via the p53-RE2, but not RE1 or RE3 (Figure 3B). We then examined if p53 binds to the *Sept4/ARTS* gene promoter at the p53-RE2 site by performing a chromatin-associated immunoprecipitation (ChIP) assay. Consistently, p53 markedly associated with the promoter fragment harboring the RE2, but not RE1 (Figures 3C and 3D). Taken together, these results demonstrate that *ARTS* is a bona fide p53 target gene.

**Figure 3.**
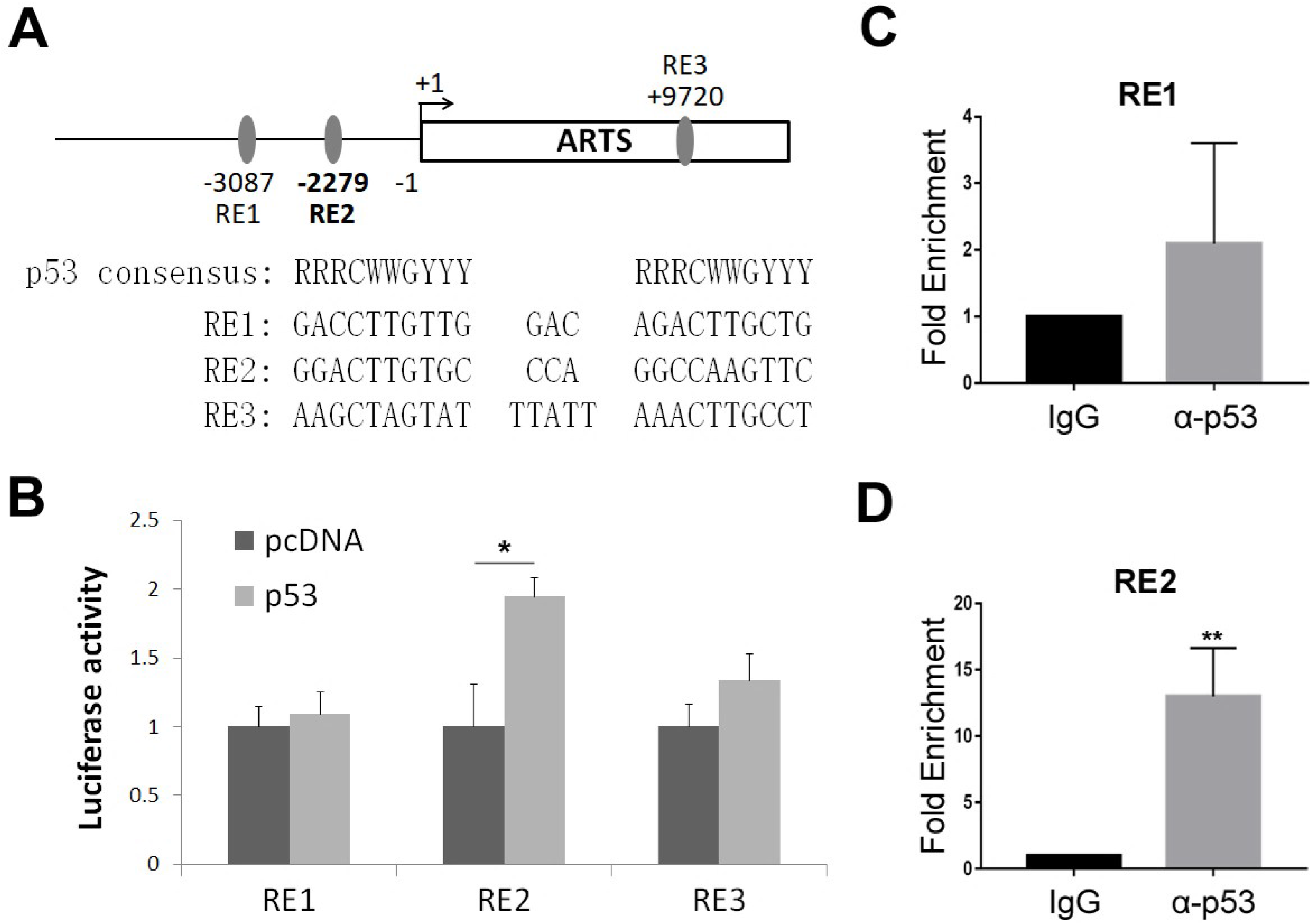
p53 associates and activates ARTS promoter. (A) A schematic of predicted p53-responsive elements on the promoter or within the intron of ARTS. (B) p53 induces the expression of a luciferase reporter gene driven by RE1. H1299 cells were transfected with plasmids as indicated in the Materials and Methods for over 30 h followed by the luciferase assay. (C, D) p53 associates with RE2, but not RE1, on the ARTS promoter. HCT116 ^p53+/+^ cells were prepared for the ChIP-qPCR analysis using the antibodies as indicated.

### ARTS binds to p53 without affecting p53 protein stability

ARTS has been documented as a key pro-apoptotic protein (Larisch et al., 2000, Edison et al., 2012) acting by targeting XIAP (Gottfried et al., 2004, Garrison et al., 2011, Edison et al., 2012) and Bcl-2 (Edison et al., 2017). Thus, we sought to explore if ARTS plays a role in p53-associated apoptotic pathway, since it is a p53-inducible gene. Intriguingly, through our recent work by screening mutant p53-interacting proteins in ovarian cancer as previously described (Chen et al., 2019), we unexpectedly found that mutant p53 may interact with a peptide (KLQDQALKE) encoded by the *SEPT4* gene through a co-immunoprecipitation (co-IP) assay coupled with mass spectrometry (MS) analysis (Figure 4A). This observation prompted us to test if ARTS binds to wild-type p53 as well, because both wild-type and mutant p53 share common binding partners in many cases, such as MDM2 and TRIM71 (Nguyen et al., 2017, Chen et al., 2019). By co-expressing exogenous p53 and Flag-ARTS in H1299 cells followed by co-IP-IB assays, we found that exogenous p53 could be co-immunoprecipitated with Flag-ARTS using an anti-Flag antibody (Figure 4B). Also, Flag-ARTS was co-immunoprecipitated with exogenous p53 using an anti-p53 antibody (Figure 4C). Furthermore, the interaction between endogenous ARTS and p53 proteins was verified by a co-IP assay using an anti-ARTS antibody in H460 cells (Figure 4D). Since some of the p53-inducible proteins, such as NGFR (Zhou et al., 2016) and PHLDB3 (Chao et al., 2016) identified by the same microarray screening as mentioned above (Liao et al., 2012), can promote p53 protein turnover by binding to the latter, we tested if ARTS can do so as well. However, transient overexpression of ARTS seemed not to affect the protein level of either exogenous or endogenous p53 in H1299 and HCT116^p53+/+^ cells (Figures 4E and 4F). Thus, these results demonstrated that ARTS interacts with p53 in cancer cells, and also suggested that ARTS may regulate p53 activity through their interaction.

**Figure 4.**
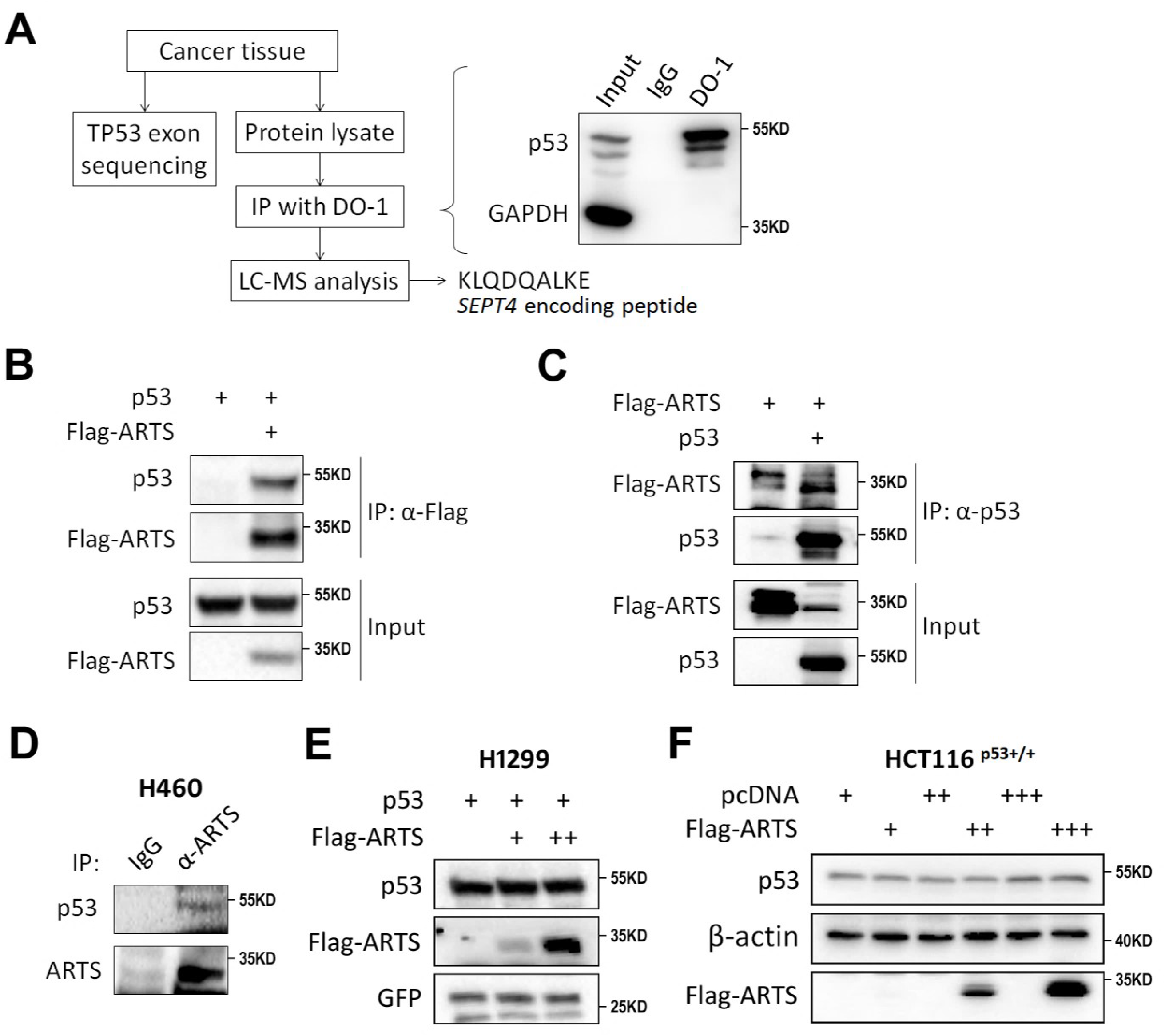
ARTS interacts with p53. (A) A schematic of screening for p53-interacting proteins in the cancer tissue harboring mutant p53-S241F. (B, C) The exogenous interactions between ARTS and p53. H1299 cells were transfected with plasmids as indicated followed by co-IP assays using the anti-flag (B) or anti-p53 (C) antibody. The bound complexes were assessed by IB analysis. (D) The endogenous interaction between ARTS and p53. The H460 cell lysates were prepared for co-IP-IB assays using the antibodies as indicated. (E, F) Ectopic ARTS does not affect exogenous (E) and endogenous (F) p53 stability. H1299 or HCT116 ^p53+/+^ cells were transfected with plasmids as indicated followed by IB analysis.

### Ectopic ARTS enhances p53-Bcl-XL interaction by sequestering p53 in the mitochondria

Although the cytosolic p53 induces apoptosis directly through the mitochondrial pathway (Green and Kroemer, 2009), the mechanism that relocates p53 to the mitochondria remains to be determined. Given that ARTS was shown to reside at the outer membrane of the mitochondria at the initiating stage of apoptosis (Edison et al., 2012), we examined if ARTS is responsible for the mitochondrial localization of p53. Through fractionation of cellular components, we observed that ectopically expressed Flag-ARTS predominantly locates at the mitochondria, and ARTS markedly increases the mitochondrial fraction of endogenous p53 (Figure 5A). The mitochondrial fraction was also used for co-IP analysis. Consistently, we verified the interaction of endogenous ARTS and p53 in the mitochondria (Figure 5B). It was noted that Doxorubicin treatment does not significantly enhance the ARTS-p53 interaction. This was probably because ARTS might recruit p53 to the mitochondria through a “hit-and-run” mechanism and it would translocate from the mitochondria to the cytoplasm and the nucleus in the later stage of apoptosis (Larisch et al., 2000). Then, we sought to determine if ARTS regulates p53’s function during mitochondrial apoptosis. In the early stage of apoptosis, p53 can binds to Bcl-XL, Bcl-2 and BAK (Green and Kroemer, 2009), we thus tested if ARTS regulates their interactions at the mitochondria. Interestingly, as shown in Figure 5C, p53 slightly bound to Bcl-XL as reported previously (Mihara et al., 2003, Moll et al., 2006), and this interaction was markedly increased when ARTS was co-expressed in cells. Therefore, these findings demonstrate that ARTS sequesters p53 in the mitochondria and consequently enhances the interaction of p53 with Bcl-XL in this subcellular compartment, and also suggest that ARTS may collaborate with p53 in triggering apoptosis, which will be addressed as follows.

**Figure 5.**
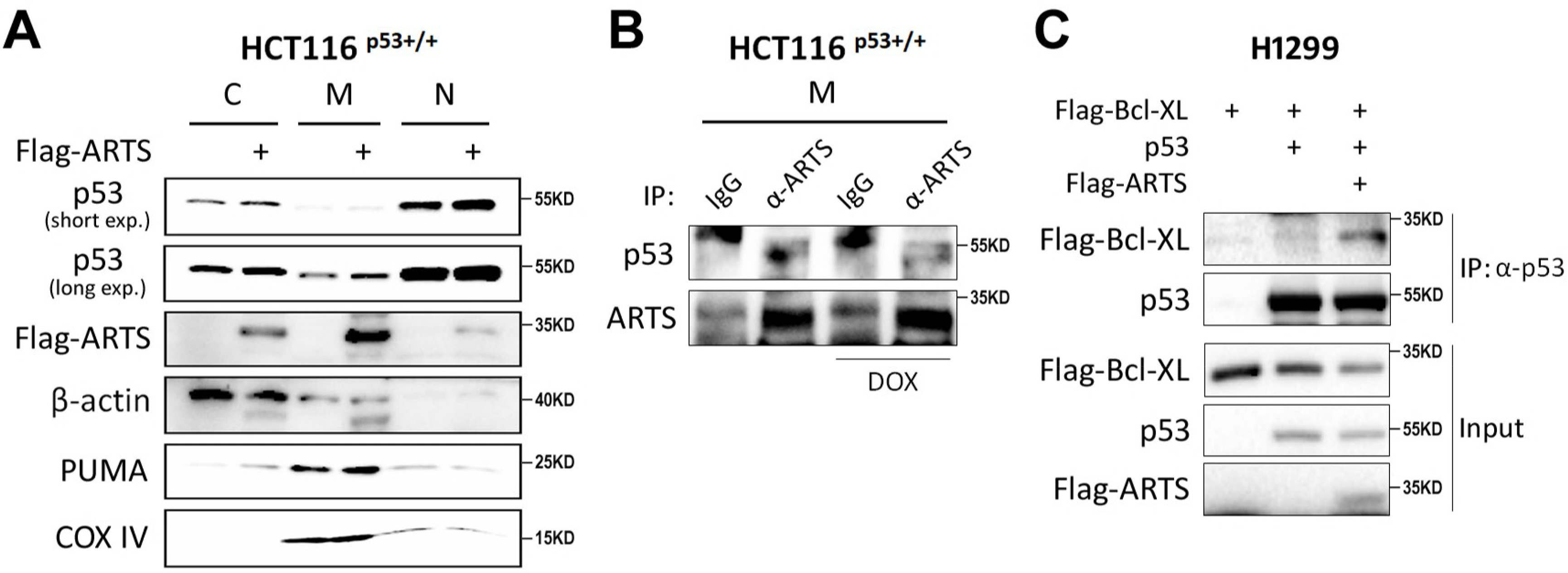
ARTS enhances p53-Bcl-XL interaction. (A) Overexpression of ARTS promotes mitochondrial accumulation of p53. HCT116 ^p53+/+^ cells were transfected with Flag-ARTS or the vector plasmid and subjected to cellular component fractionation. IB assays were performed to assess the expression of p53 and ARTS in the nucleus, cytoplasm and mitochondria, respectively. The expression of COX IV indicates the mitochondrial fraction. (B) ARTS interacts with p53 in the mitochondria. The mitochondrial fraction of HCT116 ^p53+/+^ cells was prepared for the co-IP assay using antibodies as indicated. (C) Overexpression of ARTS enhances the interaction of p53 with Bcl-XL. H1299 cells were transfected with plasmids as indicated followed by the co-IP-IB analyses.

### The p53-ARTS feedback circuit promotes apoptosis in cooperation

To determine the biological significance of the p53-ARTS feedback loop, we assessed if ARTS is involved in p53-induced apoptosis by flow cytometry analysis. Ectopic ARTS marginally induced apoptosis under the unstressed condition (Figures 6A and 6B), which is in line with the former studies (Gottfried et al., 2004, Garrison et al., 2011). Interestingly, Cisplatin or Nutlin-3 treatment significantly sensitized cancer cells to apoptosis induced by ectopic ARTS (Figures 6A and 6B), suggesting that ARTS may collaborate with activated p53 to prompt apoptosis under stress conditions. Furthermore, we examined if endogenous ARTS is required for stress-triggered apoptosis. As illustrated in Figures 6C and 6D, knockdown of ARTS significantly, though moderately, impaired DNA damage-induced apoptosis. Of note, we showed that ARTS depletion dramatically represses Nutlin-3-induced apoptosis to a greater extent than Cisplatin-induced apoptosis (Figures 6E and 6F). Considering that Cisplatin triggers apoptosis through genotoxic stress that might be partially p53-independent, while Nutlin-3 is a specific p53 agonist inducing p53-dependent apoptosis, we believed that endogenous ARTS is more selectively responsible for p53-induced apoptosis. Altogether, these results demonstrate that ARTS promotes apoptosis in cooperation with p53.

**Figure 6.**
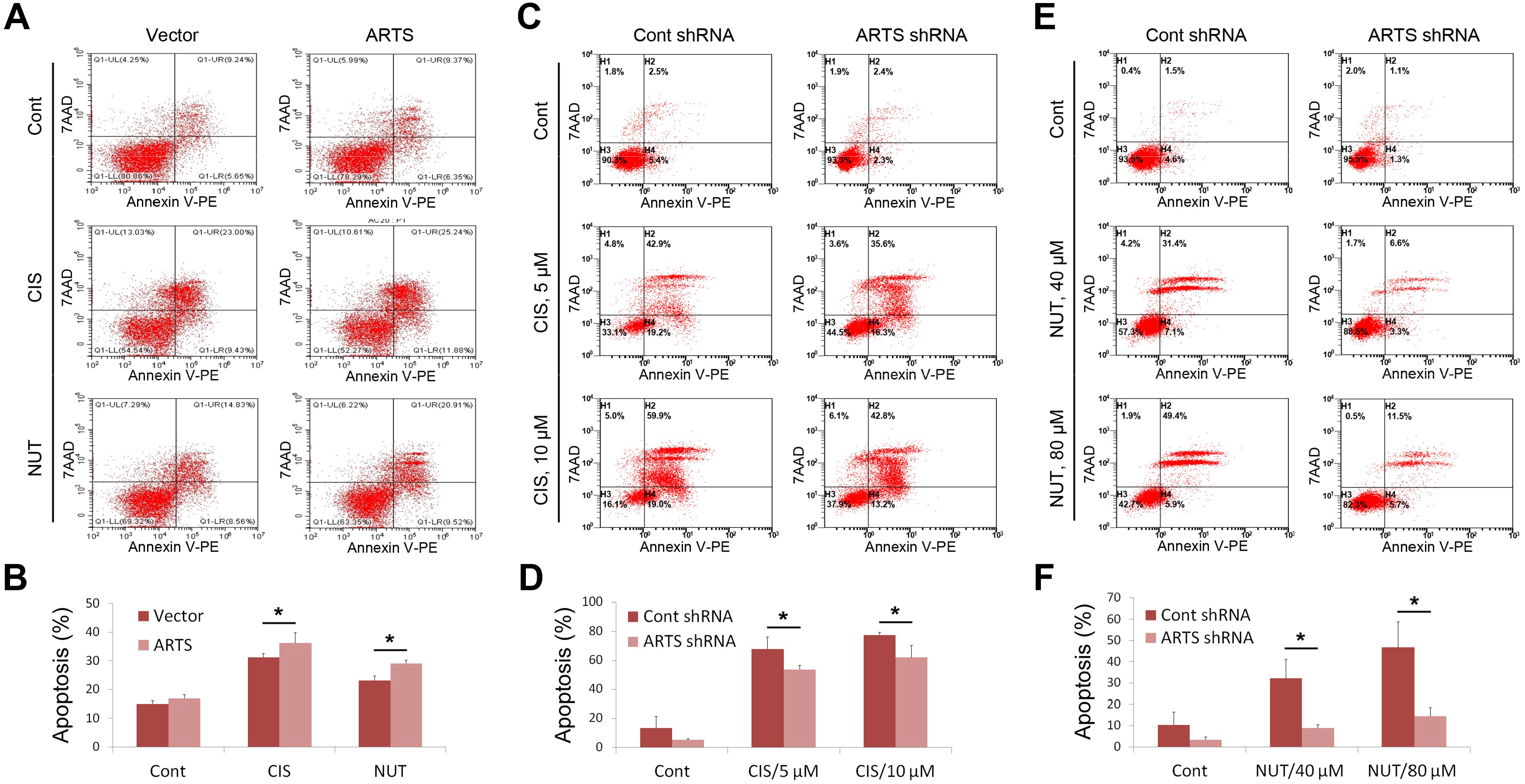
ARTS promotes apoptosis in cooperation with p53. (A, B) Overexpression of ARTS significantly augments Cisplatin- or Nutlin-3-induced apoptosis. HCT116 ^p53+/+^ cells stably overexpressing the vector or ARTS were treated with 20 μM Cisplatin or 10 μM Nutlin-3 for 48 h. The apoptosis of the treated and untreated control cells were then analyzed by flow cytometry. (C, D) Ablation of ARTS diminishes Cisplatin-induced apoptosis. H460 cells stably expressing control or ARTS shRNA were treated with 0, 5 or 10 μM Cisplatin for 48 h and subjected to flow cytometry analysis for apoptosis. (E, F) Ablation of ARTS markedly impairs p53-induced apoptosis. H460 cells stably expressing control or ARTS shRNA were treated with 0, 40 or 80 μM Nutlin-3 for 48 h and subjected to flow cytometry analysis for apoptosis.

## Discussion

The tumor suppressor p53 promotes cancer cell death through transcriptional activation of multiple pro-apoptotic genes or direct interaction with Bcl-2 family proteins in the mitochondria. Herein, we have unveiled ARTS as a novel transcriptional target and a positive feedback regulator of p53 during mitochondrial apoptosis (Figure 7). We showed that p53 transcriptionally induces ARTS expression in cancer cells and in mice (Figures 1 and 2) by binding to the ARTS promoter (Figure 3). In addition, ARTS interacts with and detains p53 in the mitochondria, resulting in increased interaction between p53 and Bcl-XL (Figures 4 and 5) and augmented apoptosis (Figure 6). Thus, our study demonstrates that ARTS plays a critical role in the p53-induced mitochondrial apoptotic pathway.

**Figure 7.**
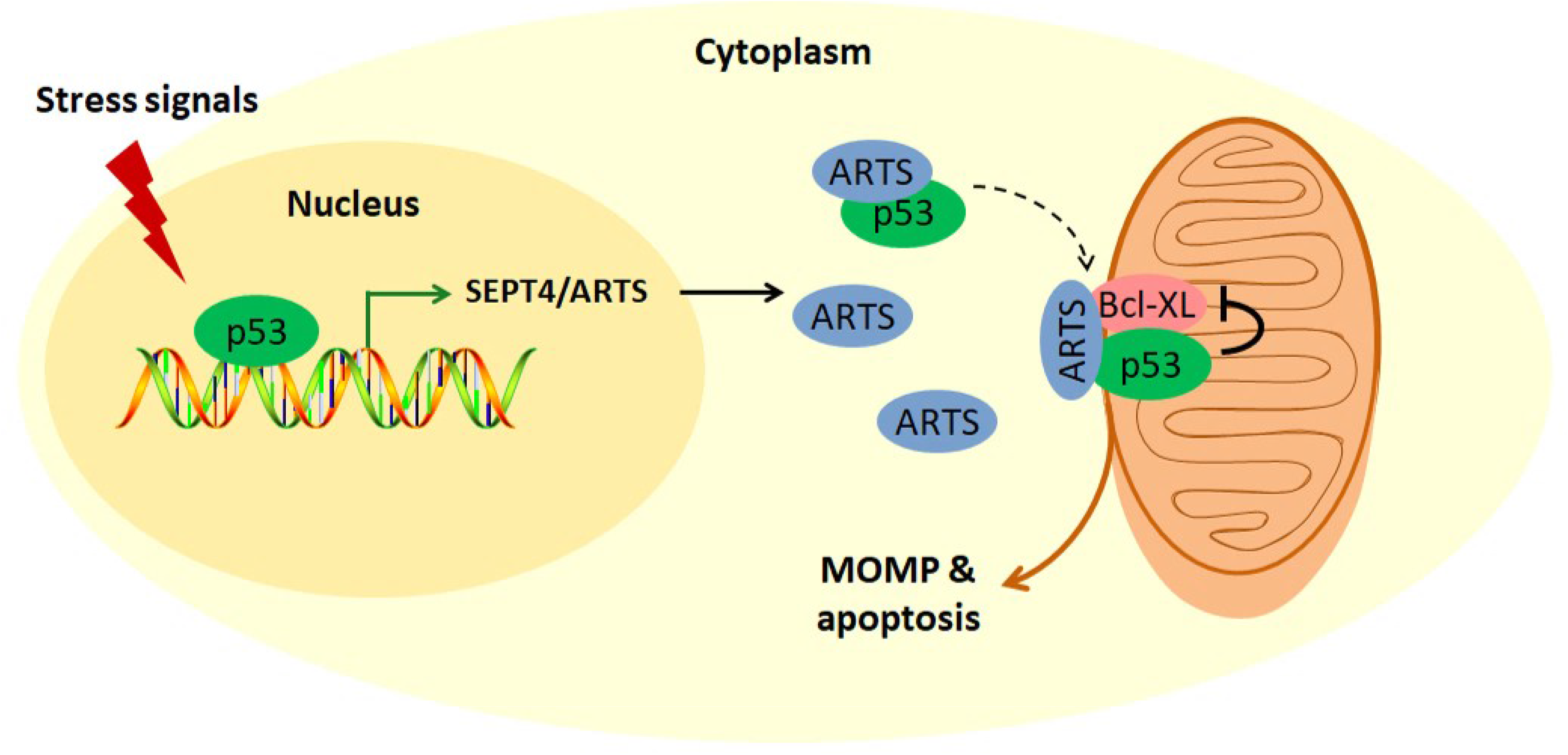
A schematic for the p53-ARTS positive feedback loop in triggering mitochondrial apoptosis. Upon various stress conditions, p53 transcriptionally induces the expression of ARTS, which in turn interacts with and relocates p53 to the mitochondria. The interaction of p53 with Bcl-XL can be enhanced by ARTS, consequently leading to MOMP and apoptosis.

To keep ARTS expression at an appropriate level is vital for cell survival, while increased expression of ARTS provokes cell death upon various stimuli. It was shown that the ARTS protein level is strictly monitored by the proteasome degradation system (Lotan et al., 2005).

The E3-ubiquitin ligase, Parkin, specifically binds to ARTS and induces its ubiquitination and degradation (Kemeny et al., 2012). Additionally, XIAP also serves as an E3-ubiquitin ligase targeting ARTS for degradation constituting a negative feedback loop (Bornstein et al., 2012). However, the mechanism for the *Sept4/ARTS* gene transcription remains unclear. Our study has revealed that DNA damage stress triggered by 5-fluorouracil, Doxorubicin or γ-irradiation elevates ARTS mRNA level in cancer cells (Figure 1E–1G, 1L) and in vivo (Figure 2), which is consistent with a previous study showing that treatment of SH-SY5Y cells with Etoposide upregulates ARTS expression (Kemeny et al., 2012). More importantly, we found that the upregulation of ARTS is dependent on p53 activation, because ectopic expression of p53 increases, while knockdown of p53 reduces, both mRNA and protein levels of ARTS (Figure 1A–1D, 1J–1L). Moreover, p53 can associate with the ARTS promoter (Figure 3D) and enhance its ability to drive the expression of a luciferase reporter gene (Figure 3B). Collectively, these findings explicitly demonstrate for the first time the mechanism underlying the *Sept4/ARTS* gene transcription by p53.

It was unexpected but interesting to notice that the *SEPT4*-encoding proteins might bind to mutant p53-Y220C through a co-IP-MS analysis in our recent study (Chen et al., 2019). The *TP53* gene is thus far the most frequently mutated tumor suppressor in human cancer. The p53-Y220C mutant was shown to not only lose the tumor suppressive activity, but also be able to boost cancer cell proliferation and migration (Chen et al., 2019). Although the function of wild-type and mutant p53s diverges in cancer development, they do usually share common interacting partners as described in previous studies (Freed-Pastor and Prives, 2012). We therefore speculated that ARTS may interact with wild-type p53 and convincingly testified this idea by showing the ARTS-p53 interaction through a set of co-IP assays (Figures 4B-D and 5B). It was further found that ARTS cooperates with p53 in bolstering mitochondrial apoptosis through improvement of the p53-Bcl-XL interaction (Figures 5C and 6). Therefore, we propose here an alternative mechanism for pro-apoptotic function of ARTS in cancer, in addition to its inhibitory effects on XIAP (Gottfried et al., 2004, Garrison et al., 2011, Edison et al., 2012) and Bcl-2 (Edison et al., 2017).

Although several p53 target genes, such as PUMA, NOXA and BAX, may be potent in executing p53’s apoptotic function, we ascertained in our study that ARTS is also involved in this process. First, although ectopic ARTS has trivial effect on apoptosis of HCT116 cells culturing in the normal condition, it significantly enhances apoptosis when cells are under stress conditions with activated p53 (Figure 6A and 6B). Additionally, depletion of endogenous ARTS significantly impairs apoptosis induced by Cisplatin or Nutlin-3 (Figure 6C–6F). Intriguingly, we found that robust expression of ARTS is more essential to Nutlin-3-induced apoptosis (Figure 6E and 6F). It is probably owing to the different molecular mechanisms behind Cisplatin- or Nutlin-3-induced apoptosis. Cisplatin triggers DNA damage response and induces apoptosis via various signaling pathways, including activation of p53 as one of the mechanisms, while Nutlin-3 selectively evokes p53-dependent apoptosis (Vassilev et al., 2004). Thus, our data strongly demonstrate that ARTS is particularly required for p53-induced apoptosis.

In conclusion, ARTS can be transcriptionally activated by p53, and endorse p53’s mitochondrial apoptotic function by binding to p53 and enhancing its interaction with Bcl-XL.

## Acknowledgements

X. Z. was supported by the National Natural Science Foundation of China (No. 81672566 and 81874053), Q. H. was supported by the National Natural Science Foundation of China (No. 81702352), and H. L. was supported by the Reynolds and Ryan Families Chair Fund of Translational Cancer. S.L was supported by Israel Science Foundation (ISF) Grant 822/12.

## Authors’ contributions

Q.H. and J.C. designed, conducted and analyzed most of the experiments; J.L. prepared the irradiated mouse tissues; Y.H. conducted the ChIP assay; S.L. provided part of reagents and helpful suggestions; S.X.Z. conducted the microarray screen and prepared part of critical materials; H.L. and X.Z. designed the experiments, analyzed the data, and composed the manuscript.

